# Comparing the performance of functional versus taxonomic metagenomics for detecting ammonia disturbances in the biogas system

**DOI:** 10.1101/2025.06.10.658511

**Authors:** Dries Boers, Olivier Chapleur, Anders F Andersson, Anna Schnürer

## Abstract

Biogas is a renewable energy source with great potential, but its production is frequently hindered by process disturbances, of which a high ammonia concentration is one common cause. It is desirable that such disturbances are found as early as possible, and metagenomics data has the potential to improve this detection. This study compares functional and taxonomic aspects of metagenomics data, hypothesizing that functional data will perform better for detecting ammonia disturbances. The hypothesis was tested by metagenomic sequencing of samples from three independent studies, which followed lab-scale reactors during ammonia disturbances. The resulting sequences were used to predict protein-coding genes, which were functionally and taxonomically annotated. The read counts of these features were fitted to disturbance states and ammonia concentrations of reactor samples using regularized regression, which allowed filtering out irrelevant features even when the number of features was much larger than the number of samples. Taxonomic data had similar or better performance in detecting ammonia disturbances and in fitting ammonia concentrations, both when analyzing separate studies as well as when analyzing the combined data of the studies. Our hypothesis that functional metagenomics would outperform taxonomic metagenomics was therefore not supported.

**Visual abstract:** 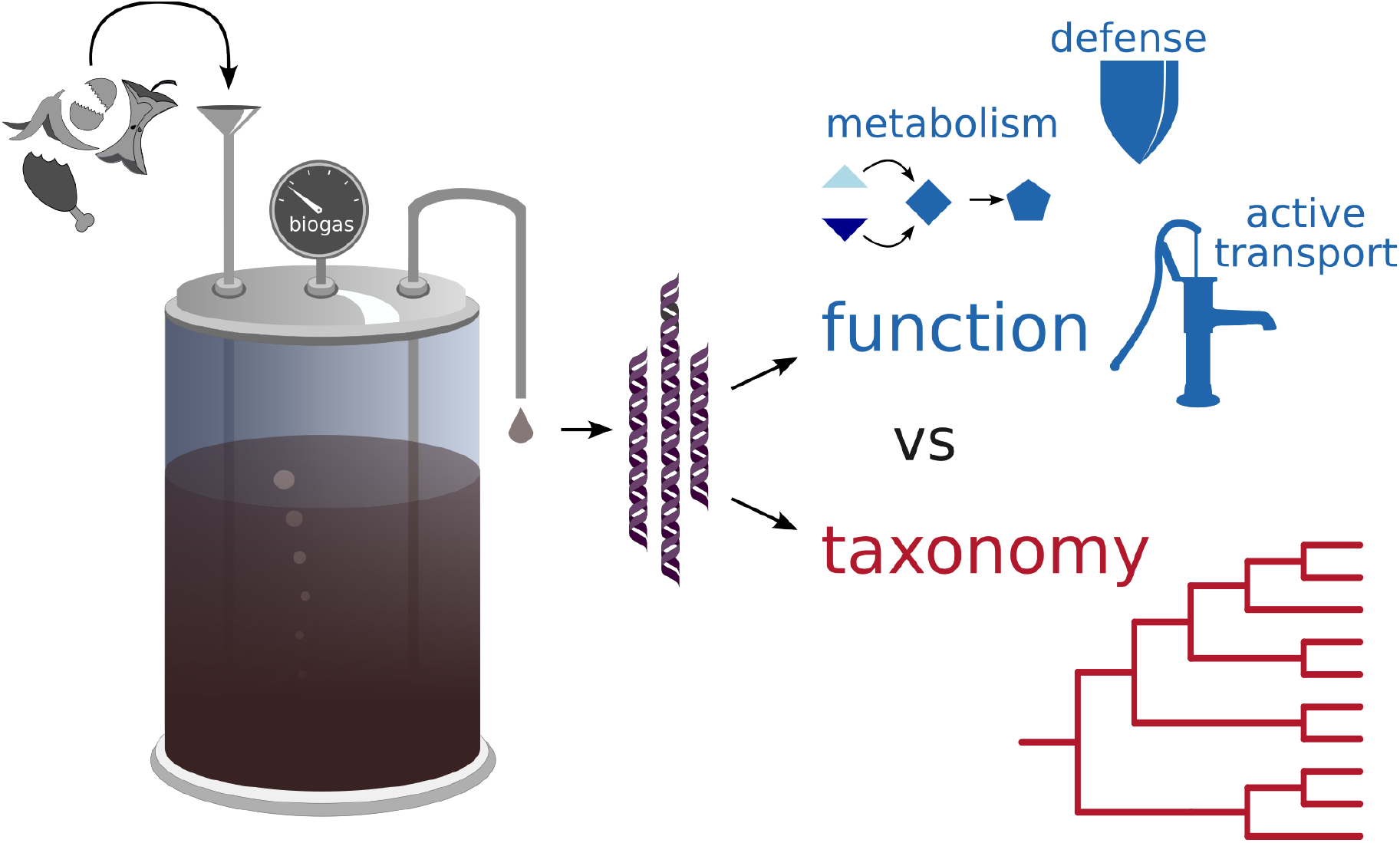

## Introduction

### The biogas system and ammonia disturbances

Biogas is an energy source which can be produced carbon-neutrally while handling society’s biodegradable waste streams. Its purified form, biomethane, can replace fossil methane gas, and anaerobic digestion, the process in which biogas is produced, allows for the recycling of the nutrients (nitrogen, phosphorous) in the input waste streams[1]. These properties make biogas an attractive energy source, and the European Union has dedicated itself to upscaling its production of biogas in the REPowerEU plan[2].

A challenge in biogas production is that the process is sensitive to process disturbances, which lead to reduced biogas production, and in severe cases even process failure[3]. Such disturbances have been shown to occur frequently, lasting from weeks up to months, with methane output decreasing by 30%[4]. As a preventive measure, most biogas plants are being operated at lower organic loading rate than needed for reaching optimal biogas production[3].

A common cause for disturbances of the anaerobic digestion process is ammonia. Anaerobic digestion consists of interlocking chains of microbial metabolism. The final chain in this network are the methanogens, methane-producing archaea, and these are particularly sensitive to ammonia poisoning[5]. The cellular mechanism of this inhibition is not clear, but the ability of ‘free’ ammonia (NH_3_) to permeate across membranes seems to be involved, because toxicity is lower at lower pH, at which more ammonia is converted to ammonium ions (NH_4_^+^), which are not freely-diffusing due to their charge[6]. When the methanogens are inhibited, this leads to the accumulation of upstream metabolites, such as hydrogen and carbon dioxide, and also acetate and other volatile fatty acids (VFAs)[7]. Such build-ups are therefore clear signals of a disturbance.

### Towards microbial community monitoring

Because of the cost of disturbances in biogas plants, the process parameters of anaerobic digestion are often intensively monitored. In a review of such monitoring, it was shown that volatile fatty acids (VFAs) and biogas composition are indicators of disturbances at an early stage[3]. However, it has been stated that including data on the microbial community could improve monitoring effectiveness[8,9], as such data might reveal disturbances at an earlier stage than only chemical parameters[9], and because models based upon chemical parameters may stop working after large microbiological shifts[10].

The state of a microbial community can be defined by analyzing its genetic sequences. Two widely-used techniques are available for this: amplicon sequencing, which targets specific taxonomic marker sequences, and whole-genome ‘shotgun’ sequencing, which randomly samples sequences from all genomes in a sample. This study mainly concerns itself with data generated by the latter technique, and refers to it as metagenomic data. After sequence processing, such data can be analysed from two perspectives. First, the taxonomic perspective views ‘who’ is present in a sample, that is, which taxa are present in an environment. Second, the functional perspective regards which genetic functions are present and which metabolic pathways are encoded. The two alternative perspectives raise the question which of them reflects the state of a microbial system best.

### Comparing the taxonomic and functional perspectives

Intuitively, the functional perspective looks more promising than the taxonomic perspective, as functions could build more fine-grained models of the biogas system. This intuition is further supported upon the concept of functional redundancy, in which taxa can be replaced by other taxa with identical functions without any effect on the system as a whole[11]. Furthermore, microbial traits can vary greatly between closely-related organisms, even within species[12]. A phenomenon that also supports the functional perspective is that traits can be transmitted to distant taxa in horizontal gene-transfer, although it has been proposed that such transfer mostly concerns simple traits such as antibiotic resistance, while complex traits such as methanogenesis are strongly coupled to taxonomy[13].

Findings of whether the functional or taxonomic perspective is superior have varied. In the general microbiological discussion, Xu *et al*.[14] showed that in the Human Microbiome Project, the difference in classification accuracy using taxonomic 16S data and metagenomic functional annotation was not significant. However, in a landmark paper of the microbial biogeography field, Louca *et al*.[15] showed that in the invariable micro-environments of rain-forest beaker plants, functional profiles predicted from 16S data were more constant than that 16S itself, which led them to suggest that functional data in their field should be “the baseline […], particularly when the ultimate focus is on ecosystem functioning”. Again in a medical context, Casimiro-Soriguer *et al*. [16] compared the use of functional and taxonomic profiles to predict colorectal cancer based upon fecal metagenomes from seven independent studies, finding that taxonomic data generally performed best. In summary, whether functional or taxonomic features give better performance is still “subject to debate”[17].

Even though many studies have investigated the microbial composition and activity in biogas systems, the functional perspective has rarely been compared to the taxonomic perspective. For example, Campanaro *et al*.[18] looked at taxonomy and function of the assembled metagenomes of different full-scale biogas plants, but focused on correlation of taxonomy to process parameters. In another study, Fischer *et al*.[19] followed inoculates in batch experiments, and in half of these ammonia was increased. However, while 16S profiles and transcriptomics were both analyzed, their information content was not compared. A study by Lin *et al*.[20] may be the most informative for the question in mind; they followed nine reactors over time to study the predictability of response to changes in feeding. They found that this change was both reflected in an altered taxonomy composition and in (functional) metatranscriptomic data. However, they did not compare which of these perspectives followed the change best.

### Aim and hypothesis

In this study, we aim to assess whether functional metagenomic data is more accurate than taxonomic data in representing the state of microbiological systems, specifically in the case of biogas reactors which are disturbed due to changing ammonia concentrations. The goal of this study is to contribute to the function versus taxonomy debate described above, both in general and in the specific context of the biogas system. Our hypothesis is that functional analysis will perform better than taxonomic analysis. To emphasize the confirmatory approach of this study, the hypothesis has been ‘preregistered’[21] in a document which was digitally signed before starting with data analysis (Supplementary Document 1).The study was further expanded to include a taxonomic perspective based upon 16S rRNA amplicon-based data. Amplicon data is widely used, and is generally less expensive to generate than shotgun sequencing data, so including a comparison of the two types is of general interest.

By testing this hypothesis within the biogas system, we intend to explore the possibility for improving its monitoring, both by reassessing the performance of the taxonomy perspective, which has been used in biogas research more frequently, and by including the functional perspective, which is less explored.

## Methods

### Sample selection and classification

Three independent studies were identified which all followed ammonia-induced disturbances over time in multiple lab-scale, mesophilic (37 °C) semi-continuous stirred-tank reactors. These studies are referred to as the Lemaigre study[10], the Cardona study[22] and the Ahrens study[23], and their characteristics are summarized in Table 1.

**Table 1.** Description of included studies.

| Name [ref.] | Number of reactors | Hydraulic retention time | Substrate | Ammonia source | pH range | Ammonia concentration maximum (FAN, g/L) | Produced methane range (L/day) |
| --- | --- | --- | --- | --- | --- | --- | --- |
| Lemaigre [10] | 3 + 1 control | 56 days | beet pulp | urea | 6.8 – 8.52<br>(6.29 – 7.641)<sup>a</sup> | 4.07<br>(0.0267)<sup>a</sup> | 0.00 – 65.29<br>(17.60 – 51.79)<sup>a</sup> |
| Cardona [22] | 6 | 25 days | food waste | ammonium-carbonate | 7.76 – 8.98 | 2.02 | 0.00 – 0.45 |
| Ahrens [23] | 5 | 53 days | manure, grass, wheat flour and bran | protein | 7.51 – 8.18 | 0.471 | 1.42 – 4.12 |

a. Values in parentheses concern control reactor.

For each of these studies, reactor performance parameters over time were analyzed; these included feeding parameters, gas production, free ammonia nitrogen (FAN) concentrations, VFA concentrations and 16S profiles. Based upon this analysis, a selection for metagenomics sequencing was made of available samples. The selected samples were also classified into ‘disturbed’ and ‘undisturbed’ classes based upon the same parameters; especially their VFA concentrations. DNA was extracted from the selected samples using protocols that differed between the studies. For the Lemaigre study, new extractions were made using the AllPrep PowerViral DNA/RNA Kit[24]. For the Cardona study, DNA was used that had been extracted for their original study using the PowerSoil DNA Isolation Kit[25] and which had been stored at -80 °C. For the Ahrens study, extractions were made using the FastDNA Spin Kit for Soil[26]. The resulting DNA extracts were sent to Eurofins Genomics for sequencing (Illumina Novaseq, paired end, >10 M reads per sample, 150 bp per read).

### Sequence processing

The resulting sequences were processed using an adapted version of the Snakemake metagenomics workflow nbis-meta[27]. The workflow started with standard read processing; that is, trimming using Trimmomatic[28], quality-checking using FastQC[29] and MultiQC[30], assembly into contigs (each sample individually) using Megahit[31], read alignment to their respective assembly using Bowtie 2[32] and duplicate removal using SAMtools[33].

The further processing of contigs is described in greater detail, as their functional and taxonomic annotation is of specific interest to this publication. Protein-coding genes were predicted within contigs by Prodigal[34]. For functional annotation, eggNOG-mapper 2[35] mapped the amino acid sequences of the predicted genes to the eggNOG database version 5[36], using DIAMOND[37] for alignment, and then used this alignment to assign eggNOG orthologous groups (eggNOG_OGs). eggNOG was used because it is the largest database for orthologs; it is automatically inferred based upon evolutionary patterns. Its size decreases the risk of missing functions of understudied micro-organisms, which have been suggested to be abundant in the biogas system[38]. Besides the assigned eggNOG_OG, KEGG orthology data[39] included from the eggNOG database was used for additional functional annotation levels.

For taxonomic annotation, predicted protein-coding gene amino acid sequences were aligned by CAT[40], also using DIAMOND, against the NCBI non-redundant protein database[41], after which the last common ancestor of the matches with a low E-value and large sequence similarity was determined based upon the NCBI taxonomy database[41]. It should be noted that the original implementation of CAT used a majority vote per contig, which is not included in this analysis; therefore, the current procedure is referred to as ‘gene-level annotation of taxonomy’ (GAT).

Finally, functional and taxonomic features were quantified by counting the reads mapping to every protein-coding gene predicted by Prodigal and summing these per feature.

To include amplicon sequencing data in the comparison, 16S rRNA sequencing counts were included from the original studies. These sequencing counts were based upon different sequencing protocols and also on different methods for inferring sequence clusters (using amplicon sequence variants or operational taxonomic units) and different taxonomy databases.

Importantly, GAT and 16S data types contain a hierarchical component, that is, organisms which differ at species rank may be the same when viewed at domain rank. A similar hierarchy holds for KEGG, because in that database, counts can be analyzed per ortholog (the lowest level), per module, or pathway level (the highest level).

### Statistical analysis

Statistical analysis of the functional and taxonomic feature counts was performed in R[42] using packages from the tidyverse[43]. See Fig. 2 for a visual summary of the analysis workflow. First, the read counts for all features were transformed per sample, using the centered log ratio from package vegan[44], to compensate for a compositionality bias[45]. The counts were then centered per feature, but they were not scaled, as larger differences in a feature should give it larger weight in subsequent analyses.

Regularized regression was then applied to the transformed features count data. Regularization makes large and complex models smaller and simpler by removing features, which is necessary when the number of features (far) outnumbers the number of samples in a dataset. The package glmnet[46] was used, and elastic net models were constructed using mixing parameter α = ½. The elastic net uses the regularization parameter ‘λ’ to apply lasso-like penalties based upon the number of variables, resulting in the removal of features that affect model performance least, and uses the same λ to apply ridge-like penalties based upon the sum of squared coefficients. The combination of the two types of penalties circumvents the problem of solution instability due to multicollinearity, which is incurred when only applying lasso penalization[47].

Two distinct types of regularized regression were applied. In logistic regularized regression, feature data was fit to the disturbed versus undisturbed classification. In linear regularized regression, feature data was fit to FAN concentrations, which are the primary cause of the disturbance of these reactors. Both of these data types had added value, because while the classification dataset can integrate multiple data types in a simple, explainable way, so-called separation can make the performance of models with more features than samples meaningless[48]. Linear regression is not hindered by the same phenomenon.

As negative measures of performance (loss functions) of the regularized models, the defaults for logistic and linear regression were used: deviance and mean squared error (MSE), respectively. Mean loss was computed for every value of λ, using leave-one-out cross-validation, and these means were plotted against the number of features of these models. These numbers of features are inversely related to λ, because as models become more regularized, their number of features decreases. Models with minimal mean loss and, secondarily, a minimal number of included features were selected from the fitted models. In linear regression, such minimal mean loss models could be selected directly, while in logistic regression, loss tends to decrease as models grow larger, and therefore the smallest models with near-minimal loss were selected manually instead. In data types with multiple hierarchical levels, these minimal loss models were used to determine and select the best-performing subtype. Superkingdom level for GAT data and domain and kingdom level for 16S data were not selected however, because their small number of features would likely lead to overfitting in regularized regression. The minimal mean loss models of the different data types, functional and taxonomic, were then compared to one another.

The above steps were first applied to the separate study datasets. Consecutively, a combined dataset was built from the feature counts of the three studies and this was submitted to the same regularized regression procedure as above. Finally, to get more insight in the regularized models, the features of the minimal loss models were extracted.

## Results

### Sample selection and classification

Reactor parameters were collected from three independent studies, in which biogas reactors encountering ammonia disturbances were followed over time. Three of these parameters are presented in Fig. 3, and the feeding parameters in Suppl. Table 1. In all three studies, concentrations of FAN changed over time in all reactors (except for the control reactor of the Lemaigre study). For the Lemaigre and Cardona studies, the changes were induced as part of their study design, while for the Ahrens study, this occurred due to high protein content in the substrate. Coincidentally with the changes in the FAN concentrations, a disturbance occurred, that is, the production of methane decreased, and VFA concentrations increased. The disturbances were followed by recovery events with methane production returning to initial values and VFA concentrations decreasing. The magnitude of VFA concentration peaks varied between the studies, but VFA accumulation was in all cases coupled to a decreased methane production, illustrating the imbalance in the microbial community metabolism. The duration of the disturbance, as predicted based on the VFA levels, varied between the studies. Expressed in hydraulic retention time (HRT), which quantifies how long a volume remains in the reactor, the Ahrens study showed the fastest recovery time (approx. 50 days, 1 HRT), while the Lemaigre and Cardona studies took longer to recover (both at least 100 days, which respectively corresponded to 2 and 4 HRTs). In the Ahrens study, the relative levels of change of the reactors’ FAN levels were also much smaller than those of the other studies, and their methane production never fully halted.

Time points for the Lemaigre and Cardona studies consisted of time points at the ends of phases of their feeding regimes as described in their publications, which generally matched VFA profiles. For the Cardona study, day 73 and 147 were also added, because in a principal component analysis based upon 16S data (Figure 4 of that study[22]), these samples fell outside the three sample clusters to which (end-of-feeding-phase time points) day 70, 98, 127 and 175 belong. For the Ahrens study, time points for metagenomic sequencing were selected based upon VFA concentration profiles.

The classification of all time points as disturbed or undisturbed (as indicated in Fig. 3) was primarily based upon VFA concentrations for all studies. Day 73 of the Cardona study and day 43 of the Ahrens study were left unclassified however, and left out of further analysis, as their status was considered unclear. The same was the case for day 239 of the Lemaigre study and 141 of the Ahrens study, because they were taken after disturbance, which neither corresponded to the undisturbed state, nor to the disturbed state.

### Supervised analysis: regularized regression

The metagenomic sequencing data was processed and annotated, resulting in functional and taxonomic count data, which was combined with taxonomic 16S amplicon count data. The different annotation variants of count data were used to build models; logistic regression models were used to predict samples’ disturbance status; linear regression models were used to estimate FAN concentrations, which was considered to be the principal cause of the disturbance.

For both types of regression analysis, each dataset was analyzed at multiple, hierarchical levels. For example, metagenomes could be annotated at different taxonomic ranks when using the GAT tool; the same was true for 16S sequences. When annotating a metagenome functionally using the KEGG database, this could be done at the lowest, single ortholog level, but also on a higher, module level. For each count type, per study, the performances in regularized logistic regression and linear regression per hierarchy level (Suppl. Fig. 1-3) were compared to determine an optimal level. The number of variants for each count type, per study (Suppl. Table 2-4) were also assessed, to confirm correct data labeling.

**Figure 1.**
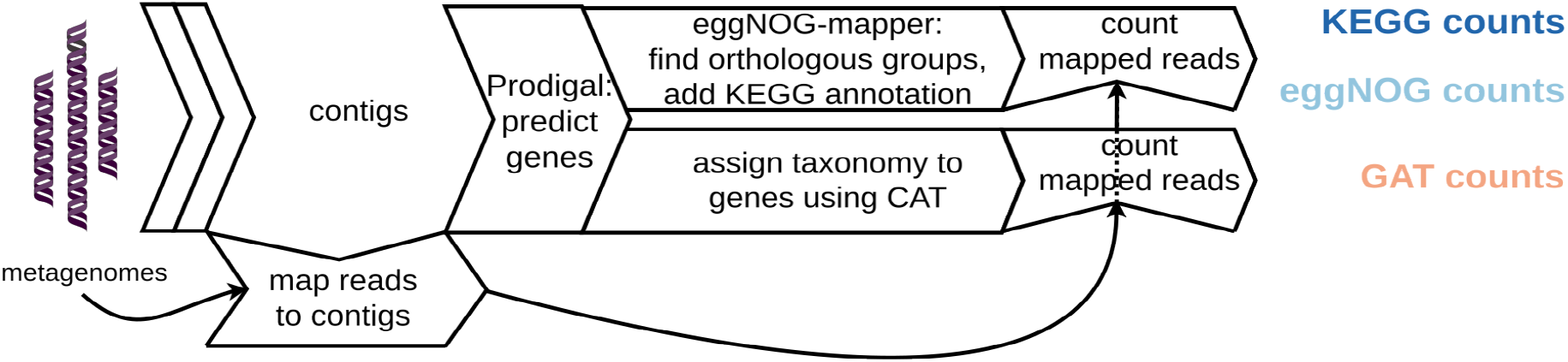
Workflow diagram of the (shotgun) metagenomics data processing to functional counts, in blue, and taxonomic counts, in light-red. GAT: gene-level annotation of taxonomy.

**Figure 2.**
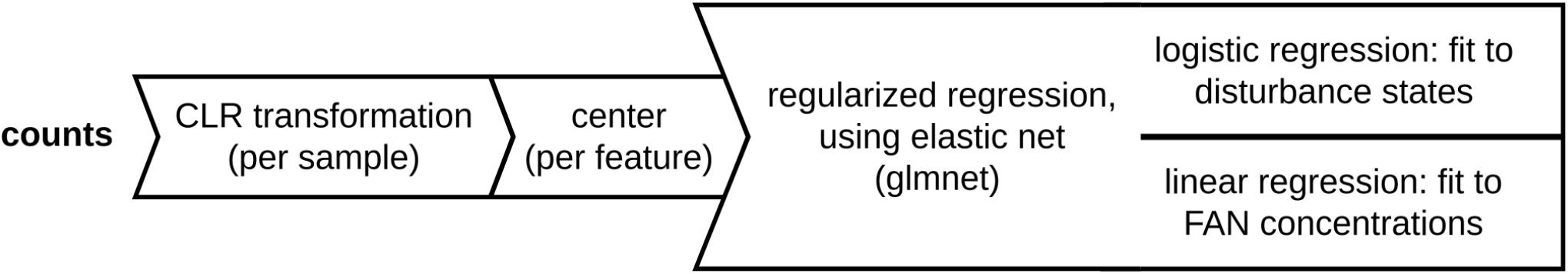
Workflow diagram of the statistical analysis of feature counts. CLR: centered log-ratio transformation. FAN: free ammonia nitrogen.

**Figure 3.**
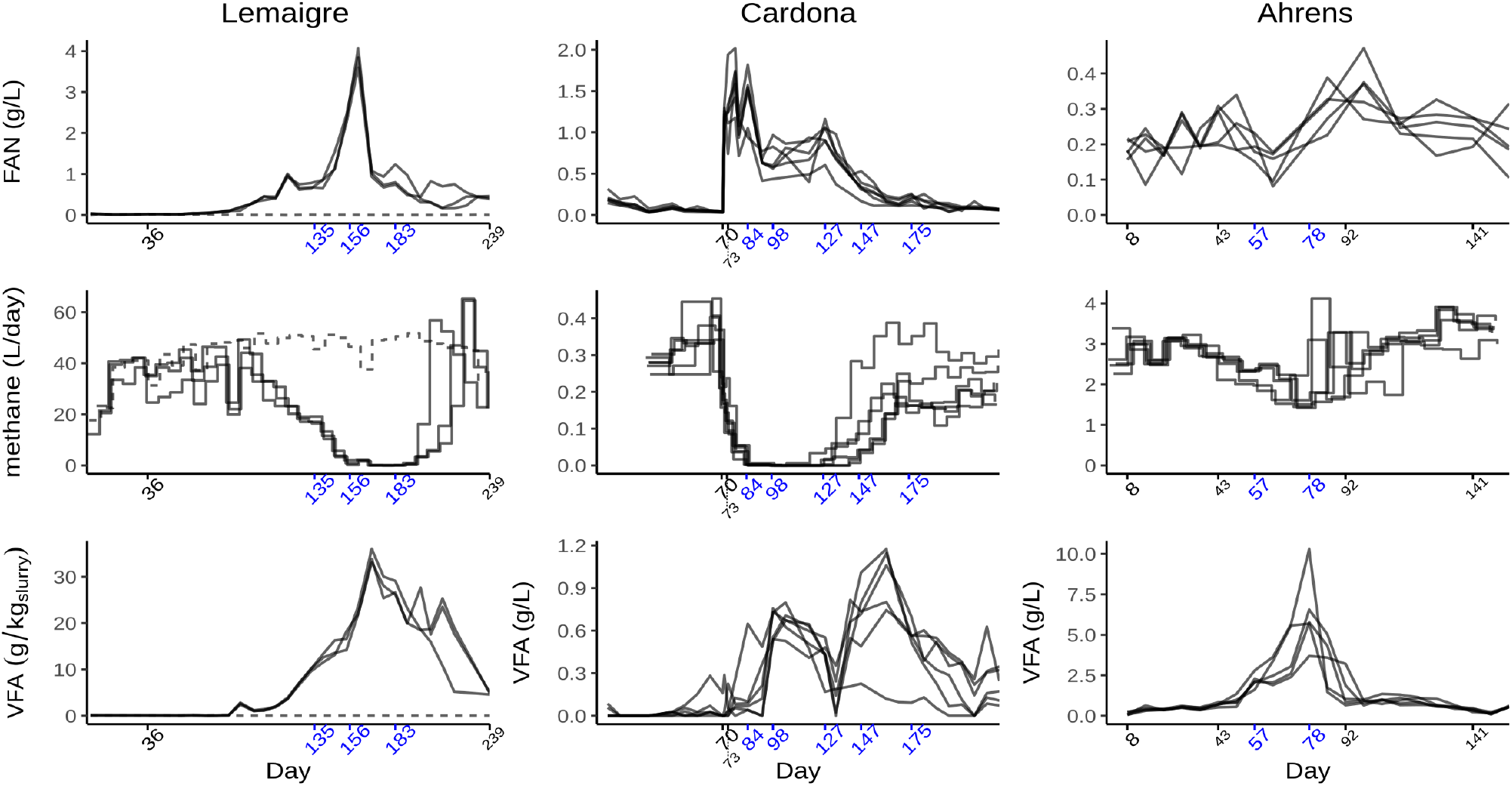
Reactor parameters for all three studies over time. In Lemaigre panels, dashed line represents control reactor without addition of extra ammonium. Breaks on time-axis indicate selected sampled time points. Blue labels indicate time points classified as disturbed, black labels with large digits indicate undisturbed time points, and black labels with small digits indicate unclassified time points. In methane panels, step graphs have been slightly shifted horizontally to prevent overlap. FAN: free ammonia nitrogen. VFA: volatile fatty acid.

**Figure 4.**
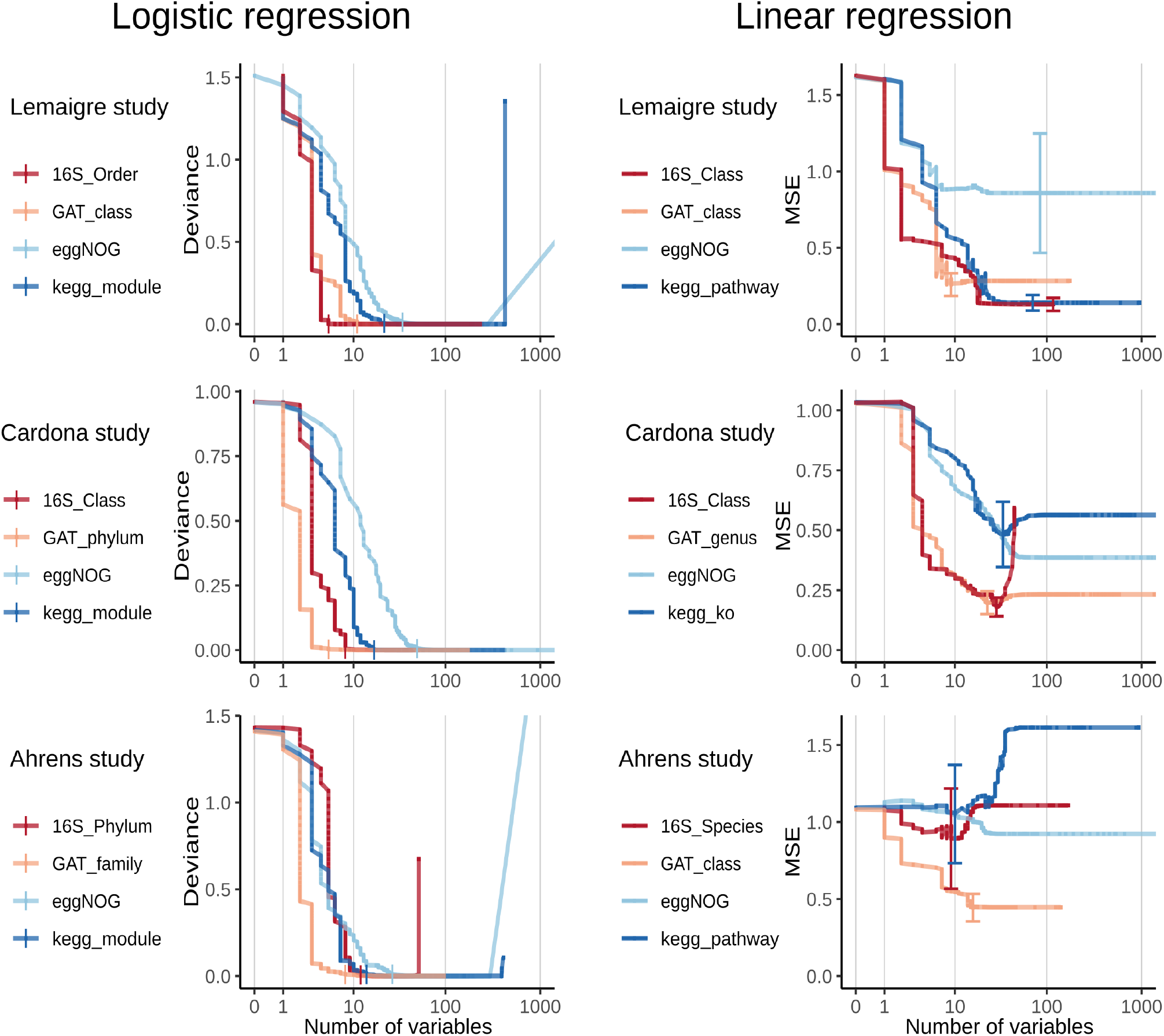
Comparison of regularized logistic and linear regression models, which are based upon functional or taxonomic count data, across studies. Models’ mean loss values, calculated by cross-validation, are plotted against the number of included variables, on a logarithmic scale. As loss function, deviance is used for logistic regression and MSE for linear regression. In logistic regression, smallest models with near-minimal mean loss (per count type, per study) are marked with a vertical bar. In linear regression, standard error bars have been added to models with minimal mean loss. MSE: mean squared error.

In the final selection of optimal hierarchy levels, the selected levels of taxonomic count types were in majority of high rank; 7/12 had rank class or higher, but low ranks such as species and genus were also selected. The selected rank was sometimes the same between logistic and linear regression; this was the case in count type GAT in the Lemaigre study and in count type 16S in the Cardona study. Sometimes the selected rank diverged substantially between logistic and linear regression however; in the Ahrens study, the selected rank for logistic regression was phylum while the rank for linear regression was species. Regarding the functional KEGG data type, the module level consistently performed best in logistic regression, while the three studies each had a different hierarchical level performing best in linear regression.

The optimal hierarchy levels for all count types are included in Fig. 4, where their logistic and linear regression performance are compared within studies. Taxonomic count types (16S and GAT) generally had better performance than the functional count types (KEGG and eggNOG), in both logistic regression and linear regression. That is, for the same number of variables, in logistic and linear regression respectively they have similar or lower mean deviance and mean MSE (both of which are loss functions, which are lower for better-fitting models). While model deviance approximated zero in all count types in logistic regression as model size increases, indicating that all count types could perfectly predict disturbance state, this low-deviance stage was reached with fewer variables by taxonomic count types. It should be noted, however, that in linear regression, in the Lemaigre study, KEGG outperformed GAT and performed similarly to 16S.

It is also noteworthy that in logistic regression, 16S outperformed GAT in the Lemaigre study, while in contrary, GAT outperformed 16S in the Cardona and Ahrens studies. Similarly, in linear regression, 16S outperformed GAT in the Lemaigre study while GAT outperformed 16S in the Ahrens study.

After separate analyses, data from the three independent studies were combined into a ‘combined study’ dataset, and analyzed in another regularized regression, which is shown in Fig. 5. While 16S data was omitted due to different indexing databases, GAT still outperformed the functional data types in logistic regression, but KEGG had the lowest MSE minimum in linear regression.

**Figure 5.**
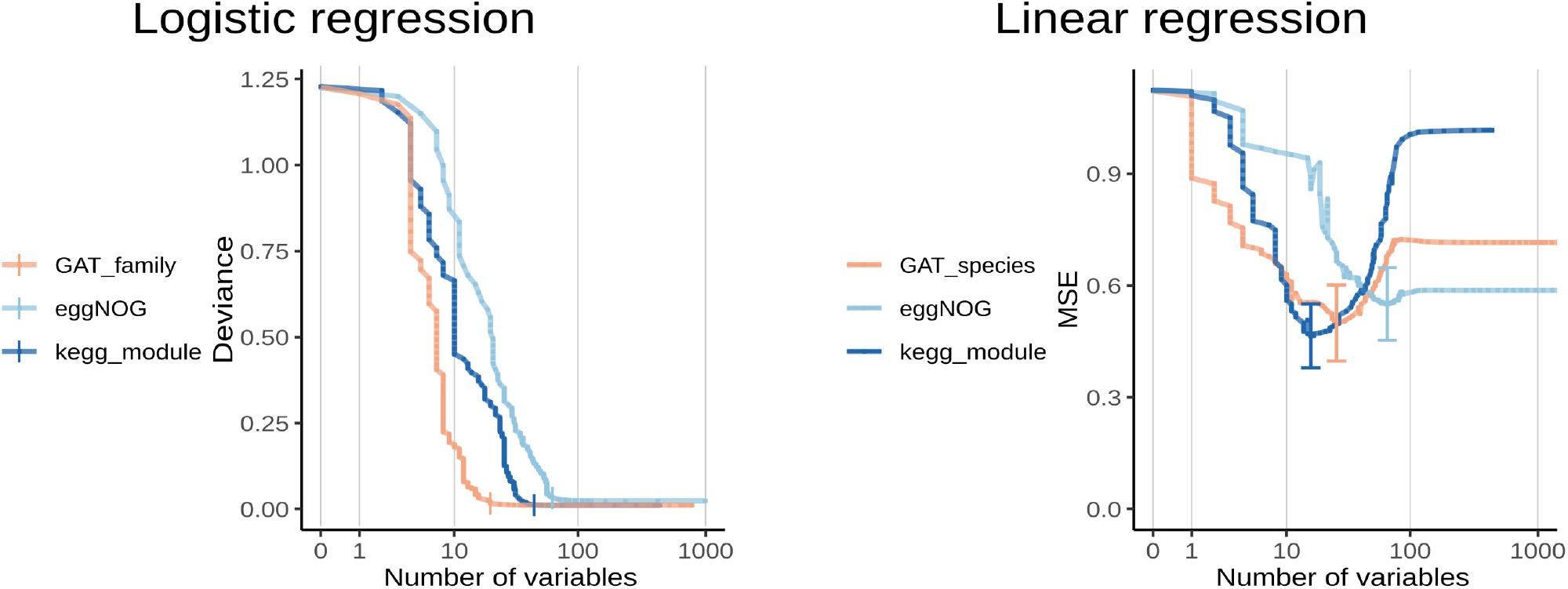
Comparison of regularized logistic and linear regression models, which are based upon functional or taxonomic count data, for combined dataset. Models’ mean loss values, calculated by cross-validation, are plotted against the number of included variables, on a logarithmic scale. As loss function, deviance is used for logistic regression and MSE for linear regression. In logistic regression, smallest models with near-minimal mean loss (per count type, per study) are marked with a vertical bar. In linear regression, standard error bars have been added to models with minimal mean loss. MSE: mean squared error.

Finally, the features of all minimal loss models were extracted, together with their regression coefficients. The number of shared features between logistic and linear regression analysis for the same feature type and study, and the number of shared features between studies for the same feature type and regression type are included in Supplementary Table 5 and Supplementary Table 6, respectively. Although an in-depth description of all included features is beyond the scope of this study, it is remarkable that in all data types, only few features are shared between selected logistic and linear regularized regression models. Only one taxon identified by GAT is shared by both logistic and linear regression, and only two of many extracted eggNOG features are shared. This was the case even when considering that count types can have different hierarchical levels for different studies. However, several taxa assigned based upon 16S data are shared, and for KEGG features, 5/61 modules were selected by both logistic and linear regression in the combined study.

An example is M00085 “Fatty acid elongation in mitochondria”. Shared features could not be straightforwardly detected in the separate studies for the KEGG data, because in that type, extracted features were from different hierarchy levels.

Also when comparing features between studies using the same regression algorithm (Supplementary Table 6), only few features are shared. While in logistic regression, for KEGG data, 30/98 features were shared between two or more studies and module M00985 “Sulfide oxidation, sulfide => sulfur” was shared between all studies, this was the only feature type where this was the case. In all other combinations of a regression type and data type, a smaller fraction of features was shared.

## Discussion

Given the interest in including microbial community data in biogas monitoring, we have compared functional and taxonomic parameters for detecting process disturbances in reactors. Our regularized regression analyses (Fig. 4 and Fig. 5) show that models based on taxonomic parameters consistently performed similarly as or better than models based on functional parameters. This is the case in three independent lab-scale studies of ammonia disturbance; both when analyzing their data separately and when analyzing the combined data. These results do not support our hypothesis that functional metagenomics data represent the biogas process better than taxonomic metagenomics data.

When comparing our results to findings reported in earlier literature, they do not match the statement by Louca *et al*.[15] that functional data should be the “baseline” of future microbial studies (they only claimed this for the context of biogeography however). Even in linear regression, taxonomic models gave a similar or better fit for FAN concentrations, which does not support the expectation that functional models, which are more fine-grained, perform better. On the other hand, our results do match the statistically insignificant difference in classification accuracy between 16S data and metagenomic annotation found by Xu *et al*.[14], as well as the findings of Casimiro-Soriguer *et al*.[16], who found taxonomic data outperforming functional data in a human microbiome context.

To explain the presented results, one could hypothesize that functional databases are even more incomplete than taxonomic databases. The biogas system includes many poorly-described micro-organisms[38]. For these, the functional annotation of orthologs is likely to be based upon experiments in distantly-related organisms, which could reduce the reliability of these annotations.

The result that between taxonomic data types, the metagenomic GAT type did not consistently outperform the 16S type is in line with recent results in human microbiome research, where metagenomic data has been found to yield “similar performances” or has performed only “slightly better”[49]. This implies that for current environmental microbiological analyses, 16S-based techniques may currently suffice; which in present are to be preferred because of their lower cost. However, the fact that different workflows, including different 16S regions and databases, have been used to annotate 16S data in different studies makes it difficult to draw conclusions here. It should also be noted, that the majority vote over contigs, which is included in the full CAT-workflow, could improve the performance of the GAT type further. This majority vote was excluded here for the sake of comparison to functional data, which always function per gene.

### Challenge: limited power

While our results do not support our hypothesis, they cannot conclusively dismiss it. Even though this study included multiple independent data sets, the number of independent reactors is still too small for a statistical test with sufficient power to compare the results of the functional and taxonomic count types. Because of the same reason, no separate dataset was used to perform an independent test of the fitted glmnet models, and to test their performance. Nonetheless, both separate and combined analyses support a difference, albeit not statistically significant. These findings can be used for consecutive studies of functional and taxonomic metagenomics data in microbiology, particularly in the biogas system.

The limited number of independent reactors is also the reason why elastic net was chosen as algorithm for supervised analysis. This approach does not allow for the modeling of non-linear effects, and while this is a limitation, it decreases the risk of overfitting the dataset.

Another factor that contributed to the risk of overfitting was the multiple hierarchy levels for each count type, further increasing the number of features (compared to the limited number of samples). A connected observation is that within a study, a non-optimal subtype of a taxonomic dataset can have worse performance than a functional subtype. For example, in the Ahrens study, 16S subtype ‘order’ is outperformed by KEGG optimal subtype ‘module’.

A final limitation related to machine learning was that multiple samples were derived from identical reactors, violating the generalized linear model assumption of independent observations. The risk of the conclusions in this study being affected is decreased by the fact that these limitations apply to both types of analyses, functional and taxonomic.

The lack of shared features selected by glmnet within count types, both between regression types for the same study and between studies for the same regression type, puts the explainability of the models into question. Furthermore, KEGG is the only count type in which a larger fraction of features was shared, between studies for the same regression type. This could support the notion that while functional metagenomics does not outperform taxonomic metagenomics, the former includes more consistent feature profiles and is therefore more explainable. Confirmation of this observation through follow-up research is needed.

### Perspectives and implications

To follow up on this research the performance of other data types could be assessed. Transcriptomic sequencing data would certainly be of interest, as it might give a more reliable functional perspective than metagenomic data, showing which genes are actively being transcribed, and potentially translated, to functional proteins. While the potential of metatranscriptomics is great, handling RNA is more demanding than both 16S and metagenomics. Implementing RNA-based monitoring in biogas processes would therefore likely be more challenging than implementing DNA-based monitoring. Other possible workflows for generating sequencing count data include assembly-free annotation tools, or 16S-based functional annotation tools. We do not, however, expect such changes in the sequence processing to alter the conclusions of our study.

The implications of the functional metagenomics not outperforming taxonomic metagenomics, as well as taxonomic metagenomics not consistently outperforming 16S is that in monitoring of the biogas system, metagenomic sequencing might have little added value compared to amplicon sequencing, which is cheaper. Functional analysis could still be of value, however, when trying to understand the mechanisms of changes in a microbial system.

## Acknowledgments

The authors would like to thank Sébastien Lemaigre, Xavier Goux and Magdalena Calusinska for sharing samples from the Lemaigre study.

Dries Boers would like to thank the Swedish Bioinformatics Advisory Program, and his program advisors John Sund and Lokeshwaran Manoharan in particular. He would also like to thank Reza Belaghi for advice on classification algorithms; Stan van Lier for valuable discussions on performance metrics in regularized regression; and Nils Weng, Jonas Ohlsson and Malin Tiefensee for helpful comments on the manuscript.

This work was supported by the Swedish University of Agricultural Sciences (SLU) and the Swedish Energy Agency, project no. P2022-00552. High-throughput sequence processing was enabled by resources provided by the National Academic Infrastructure for Supercomputing in Sweden (NAISS), partially funded by the Swedish Research Council through grant agreement no. 2022-06725.

