## Supplementary Material for "Comparing the performance of functional versus taxonomic metagenomics for detecting ammonia disturbances in the biogas system"

Supplementary Document 1. Preregistered hypothesis. See next page. A digitally signed version could not be included in the current supplementary material format, but is available upon request.

Page: **Hypotheses and predictions**

Created by: Dries Boers (2022-09-09 11:37:41)

Modified by: Dries Boers (2022-09-09 12:09:48)

ELN-P-3073-

### Content

The hypotheses and predictions below were defined before sequencing datasets from LIST institute and Chapleur group were downloaded from Eurofins servers.

### Content

Hypothesis 1: Functional metagenomics represents the biogas process better than taxonomic metagenomics.

Prediction 1.1: When comparing different replicate reactors for a timepoint, functional metagenomic data will have a smaller variance compared to taxonomic metagenomic data.

Supplementary Table 1. Feeding phases for Cardona and Lemaigre studies.

| | Feed type | Day 1-70 | Day 70-84 | Day 84-129[a] | Day 129-231 | |
|---|---|---|---|---|---|---|
| Cardona study | Food biowaste (gCOD/L/day) | 0.5 | 0.5 | 0 | 0.5 | |
| | Ammonia (g/L/day) | 0 | 4[b] | 0 | 2 | |
| | **Feed type** | **Week 1-8** | **Week 8-19** | **Week 20-22** | **Week 23-26** | **Week 27-34** |
| Lemaigre study[c,d] | Carbon: beet pulp (gVS/L/week) | 10-13[e] | 10-13 | 10-13 | 0 | 8-13[e] |
| | Nitrogen: urea (g/L/week) | 0 | 1-4[f] | 8-10[f] | 0 | 0 |

a. Feeding and reactor flowthrough was fully stopped during this phase.

b. Ammonia was added on day 70 to reach a concentration of 4 g/L for all reactors.

c. Feeding varied per week within ranges.

d. Control reactor was fed constantly with 8-13 gVS/L/week beet pulp.

e. Feeding was built from lower values in first two weeks of phase.

f. Urea feeding was gradually increased during phases.

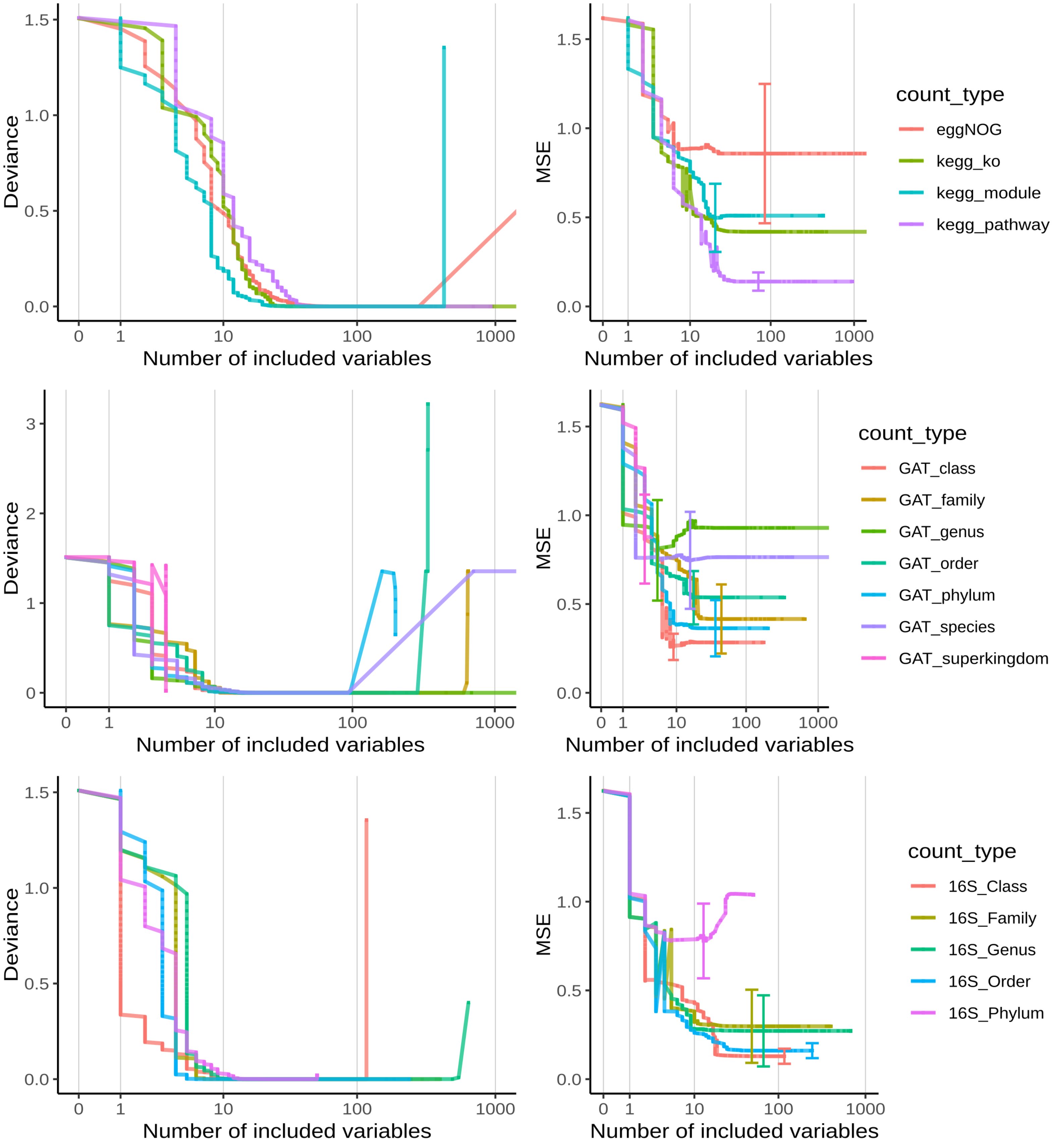


Supplementary Figure 1. Selection of optimal hierarchy level per count type. Here based upon the Lemaigre study. Count types with low loss scores with a small number of included variables are selected. Left panels are based upon logistic regression, right panels are based upon linear regression.

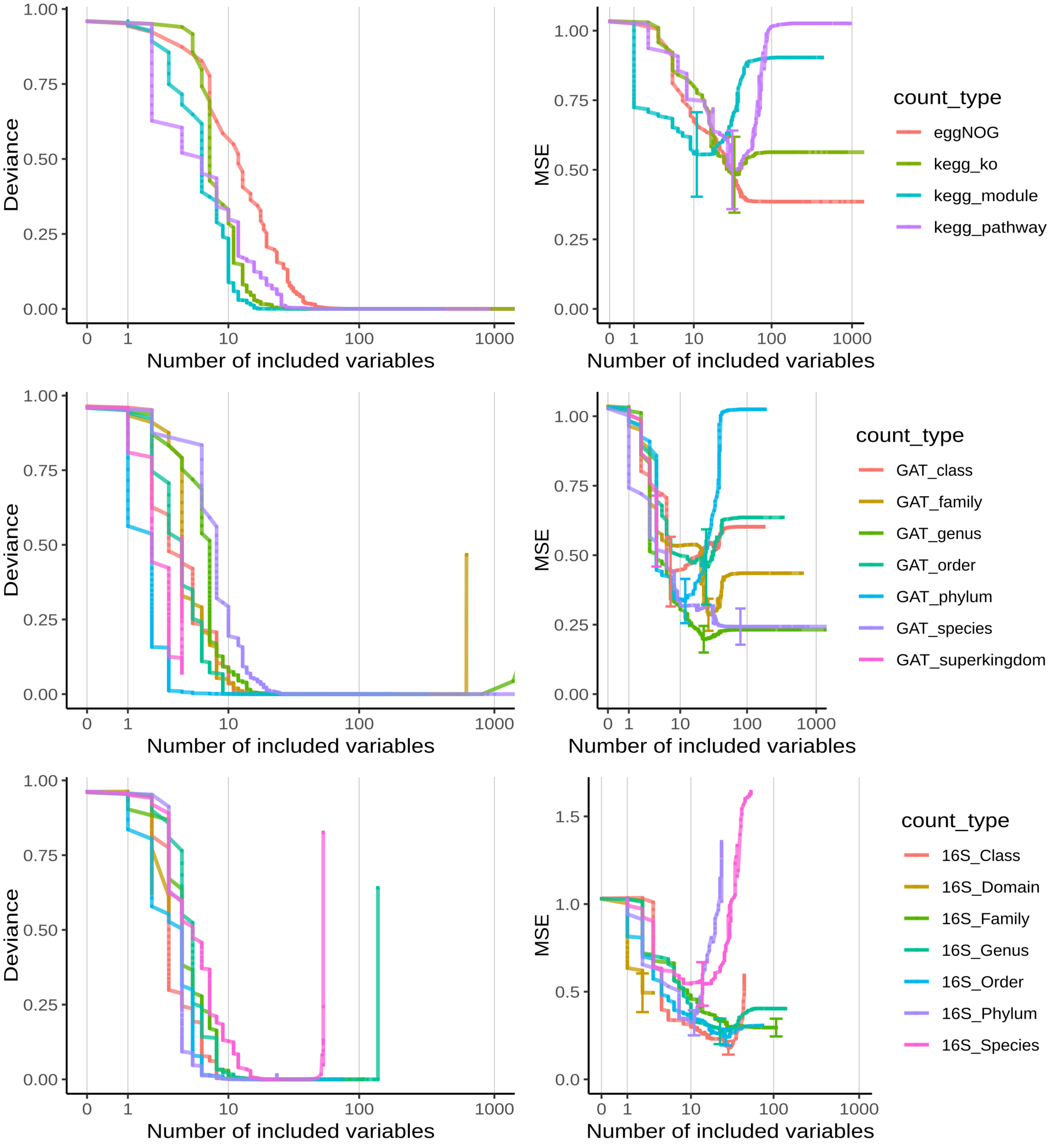


Supplementary Figure 2. Selection of optimal hierarchy level per count type. Here based upon the Cardona study. Count types with low loss scores with a small number of included variables are selected. Left panels are based upon logistic regression, right panels are based upon linear regression.

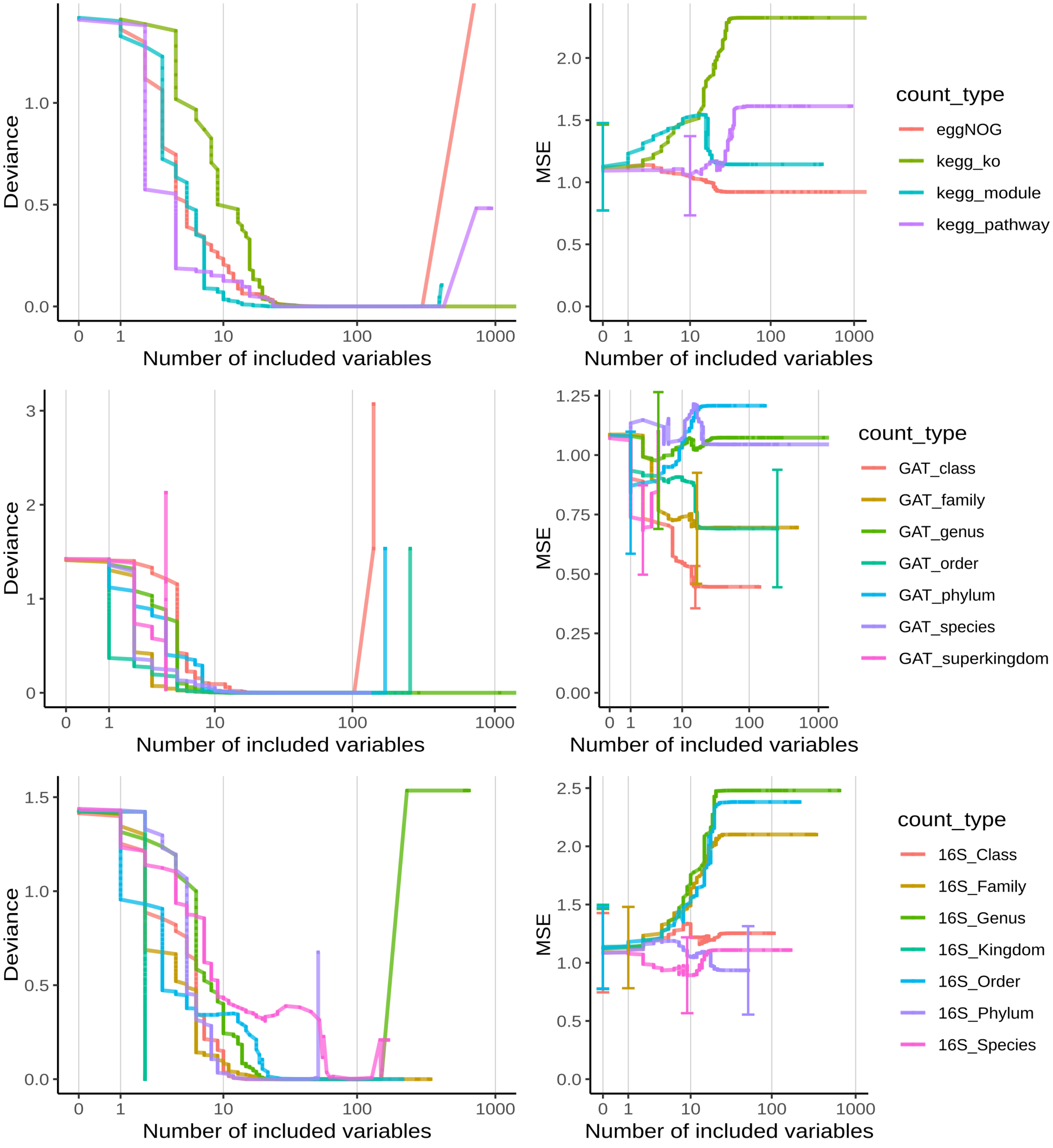


Supplementary Figure 3. Selection of optimal hierarchy level per count type. Here based upon the Ahrens study. Count types with low loss scores with a small number of included variables are selected. Left panels are based upon logistic regression, right panels are based upon linear regression.

Supplementary Table 2. Number of functional metagenomic annotation variants, per hierarchical information level and per study.

| | KEGG Pathways | KEGG Modules | KEGG ko | eggNOG |
|---|---|---|---|---|
| **Lemaigre study** | 952 | 426 | 9534 | 489070 |
| **Cardona study** | 936 | 424 | 9647 | 486519 |
| **Ahrens study** | 934 | 411 | 8180 | 329520 |

Supplementary Table 3. Number of gene-level annotation of taxonomy (GAT) variants, per taxonomic rank and per study. While it is unexpected that the unique number of phyla is larger than the unique number of classes, this is due to a large number of genes not being further annotated by GAT than the phylum rank (data not shown).

| | Superkingdom | Phylum | Class | Order | Family | Genus | Species |
|---|---|---|---|---|---|---|---|
| **Lemaigre study** | 4 | 201 | 175 | 340 | 652 | 2575 | 9775 |
| **Cardona study** | 4 | 181 | 174 | 330 | 625 | 2261 | 7778 |
| **Ahrens study** | 4 | 170 | 141 | 255 | 489 | 1786 | 6201 |

Supplementary Table 4. Number of 16S annotation variants, per taxonomic level and per study.

| | Domain | Kingdom | Phylum | Class | Order | Family | Genus | Species |
|---|---|---|---|---|---|---|---|---|
| **Lemaigre study** | *NA* | *NA* | 51 | 117 | 245 | 400 | 677 | *NA* |
| **Cardona study** | 3 | *NA* | 24 | 45 | 74 | 107 | 138 | 54 |
| **Ahrens study** | *NA* | 2 | 52 | 106 | 215 | 342 | 645 | 169 |

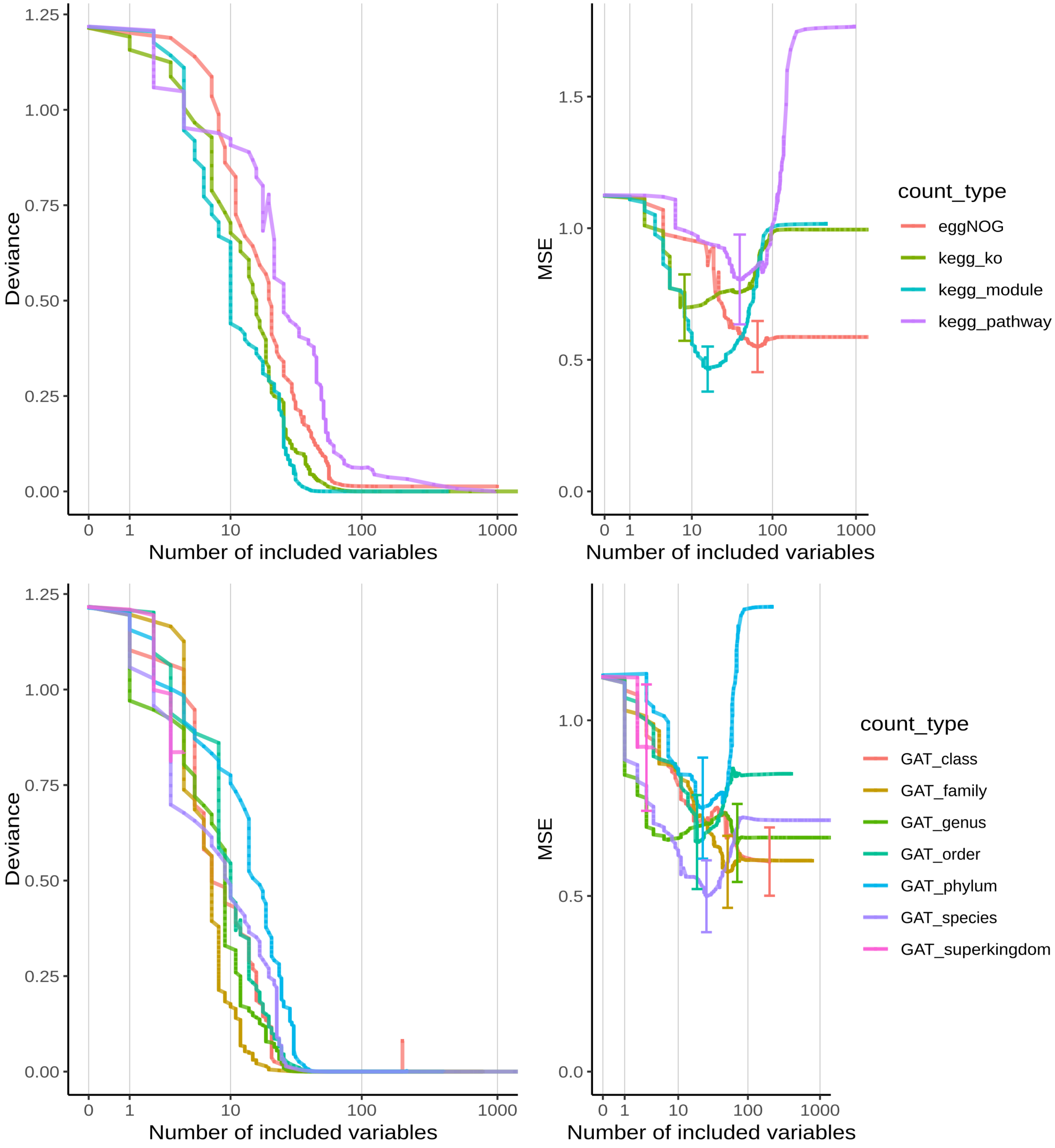


Supplementary Figure 4. Selection of optimal hierarchy level per count type. Here based upon the combined analysis. Count types with low loss scores with a small number of included variables are selected. Left panels are based upon logistic regression, right panels are based upon linear regression.

Supplementary Table 5. Number of shared extracted features beween logistic and linear regression, per count type and study.

| | KEGG | eggNOG | GAT | 16S |
|---|---|---|---|---|
| **Lemaigre study** | 0/93 | 0/120 | 0/20 | 6/122 |
| **Cardona study** | 0/51 | 1/348 | 1/28 | 6/37 |
| **Ahrens study** | 0/24 | 1/107 | 0/24 | 5/21 |
| **combined study** | 5/61 | 0/129 | 0/46 | NA |

Supplementary Table 6. Number of shared extracted features beween all studies, per count type and regression type.

| | KEGG | eggNOG | CAT | 16S |
|---|---|---|---|---|
| **Logistic regression** | 30/98 | 14/175 | 9/44 | 4/25 |
| **Linear regression** | 6/131 | 20/529 | 5/74 | 5/155 |
